# Cytoskeletal association of ATP citrate lyase controls the mechanodynamics of macropinocytosis

**DOI:** 10.1101/2022.10.05.511007

**Authors:** Joseph Puccini, Dafna Bar-Sagi

**Author notes:** Correspondence (D. Bar-Sagi).

## Abstract

Macropinocytosis is an actin-dependent mode of non-selective endocytosis that mediates the uptake of extracellular fluid-phase cargoes. It is now well-recognized that tumor cells exploit macropinocytosis to internalize macromolecules that can be catabolized and used to support cell growth and proliferation under nutrient limiting conditions. Therefore, the identification of molecular mechanisms that control macropinocytosis is fundamental to the understanding of the metabolic adaptive landscape of tumor cells. Here we report that the acetyl-CoA producing enzyme, ATP citrate lyase (ACLY), is a key regulator of macropinocytosis and describe a heretofore unappreciated association of ACLY with the actin cytoskeleton. The cytoskeletal tethering of ACLY is required for the spatially-defined acetylation of heterodimeric actin capping protein, which we identify as an essential mediator of the actin remodeling events that drive membrane ruffling and macropinocytosis. Furthermore, we identify a requirement for mitochondrial-derived citrate, an ACLY substrate, for macropinocytosis, and show that mitochondria traffic to cell periphery regions juxtaposed to plasma membrane ruffles. Collectively these findings establish a new mode of metabolite compartmentalization that supports the spatiotemporal modulation of membrane-cytoskeletal interactions required for macropinocytosis by coupling regional acetyl-CoA availability with dynamic protein acetylation.

**Significance:** The scavenging of extracellular macromolecules via macropinocytosis is a key adaptive mechanism that supports the metabolic fitness of cancer cells. Although the importance of macropinocytosis for sustaining tumor cell growth under nutrient limiting conditions is well documented, less is known about the molecular mechanisms by which macropinocytosis is regulated. This study describes a previously uncharacterized dependence of macropinocytosis on the compartmentalized generation of acetyl-CoA through the association of ACLY with the actin cytoskeleton. This metabolic channeling process establishes a new mechanistic framework for understanding actin remodeling events that drive macropinocytosis.

## Introduction

Engagement of nutrient scavenging pathways by oncogenic signaling is a key adaptive strategy that confers metabolic flexibility to cancer cells (1). This has been well described in the context of oncogenic RAS-expressing cancer cells, which upregulate macropinocytosis, a mechanism of non-selective, fluid-phase endocytosis (2–5). Macropinocytosis is initiated by the formation of membrane protrusions called membrane ruffles, a process dependent on the coordinated polymerization, crosslinking and branching of cortical filamentous actin (F-actin) (6). As these membrane protrusions extend outwards and collapse back onto the plasma membrane, they envelop extracellular fluid-phase cargoes into endocytic vesicles called macropinosomes (7). Cancer cells use macropinocytosis to scavenge extracellular macromolecules and necrotic cell debris, that are subsequently catabolized in the lysosome to yield anabolic substrates that fuel cell growth and proliferation (7). The importance of this nutrient scavenging pathway for the metabolic fitness of cancer cells is evidenced by studies demonstrating that genetic or pharmacological blockade of macropinocytosis in vivo attenuates tumorigenesis (3–5, 8). As such, there is a growing interest in defining and characterizing the key regulatory steps that control this process. Although progress has been made towards the identification of effectors and regulators of macropinocytosis (4, 5, 8–12), the interdependence between cellular bioenergetics and macropinocytosis remains poorly understood.

In this study we demonstrate that macropinocytosis is dependent on ATP citrate lyase (ACLY), an enzyme that catalyzes the conversion of mitochondrial-derived citrate and co-enzyme A (CoA) to oxaloacetate and acetyl-CoA (13). We provide evidence that ACLY is an F-actin associated protein and describe a link between ACLY-mediated production of acetyl-CoA and the regulation of actin cytoskeleton dynamics that are required for membrane ruffle formation. The association between F-actin and ACLY defines a new mechanism for the compartmentalized production of acetyl-CoA, and establishes a functional interdependence between specific intracellular metabolic intermediates and nutrient supply via macropinocytosis.

## Results

### ACLY activity is required for macropinocytosis

To explore the relationship between macropinocytosis and cellular bioenergetics, we have assessed the effects on macropinocytosis of acute pharmacological modulation of biosynthetic pathways that are critical for cancer cell fitness. This analysis revealed that treatment of cells with the ACLY inhibitor BMS-303141 (14, 15), caused a marked reduction in the uptake of dextran (a selective marker of macropinocytosis) in a panel of macropinocytic RAS mutant cell lines (Fig. 1A, B). A similar inhibitory effect was displayed by a structurally unrelated, allosteric inhibitor of ACLY, NDI-091143 (16), indicating that ACLY activity might be required for macropinocytosis (Fig. 1C, D). While the role of ACLY-dependent acetyl-CoA production in promoting lipogenesis and maintaining global histone acetylation is well established (17–20), a link between ACLY activity and the regulation of endocytic processes, such as macropinocytosis, has not been described. We therefore sought to further validate this link using a genetic approach involving the generation of ACLY knockout (KO) clones of T24 cells by CRISPR/Cas9 genome editing (Fig. S1A-C). Previous reports have demonstrated that acyl-CoA synthetase short chain family member 2 (ACSS2) is able to compensate for genetic deletion of ACLY by synthesizing acetyl-CoA from acetate (15, 21) (Fig. 1E). Therefore, we assessed the macropinocytic capacity of ACLY WT and KO clones in the presence and absence of exogenous acetate. In contrast to ACLY WT clones, which displayed comparable levels of dextran uptake independently of acetate, ACLY KO clones showed impaired dextran uptake following acetate withdrawal (Fig. 1F-H). Additionally, re-expression of WT ACLY, but not catalytically inactive ACLY (H760A) (15), was sufficient to restore dextran uptake in ACLY KO cells, rendering them insensitive to acetate withdrawal (Fig. 1I-K). Taken together, these results uncover a previously uncharacterized dependence of macropinocytosis on ACLY activity and acetyl-CoA production.

**Figure 1.**
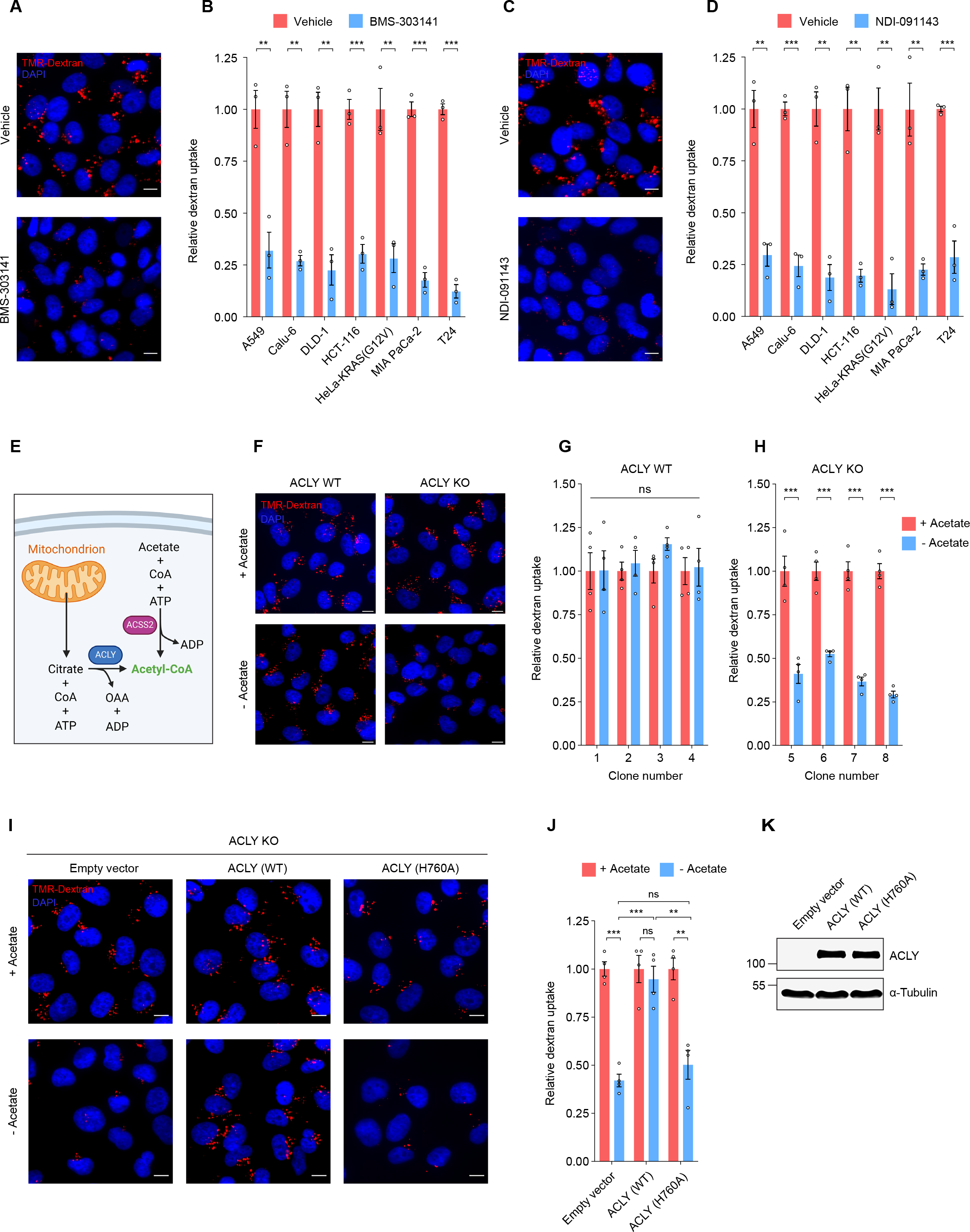
ACLY activity regulates macropinocytosis. **(A**, **B)** Fluorescence micrographs and quantification of TMR-dextran uptake in cell lines treated with ACLY inhibitors (1 hour) BMS-303141 and **(C**, **D)** NDI-091143. **(E)** Schematic showing the major intracellular sources of acetyl-CoA. ACLY uses mitochondrial-derived citrate and coenzyme A (CoA) to synthesize acetyl-CoA and oxaloacetate (OAA). ACSS2 utilizes acetate and CoA to synthesize acetyl-CoA. **(F)** Fluorescence micrographs and quantification of TMR-dextran uptake in **(G)** ACLY WT and (**H**) ACY KO T24 clones in the presence or absence of sodium acetate. **(I)** Fluorescence micrographs and **(J)** quantification of TMR-dextran uptake in ACLY KO T24 cells reconstituted with empty vector, WT ACLY or H760A (catalytically inactive mutant) ACLY in the presence or absence of sodium acetate. **(K)** Immunoblot showing ACLY expression in ACLY KO T24 cells reconstituted with empty vector, WT ACLY or H760A ACLY. α-tubulin was used as a loading control. All images and immunoblots are representative. Scale bar for all images = 10 μm. For TMR-dextran uptake assays, at least 500 cells were counted per biological replicate (n=3-4). Data represented as mean ± s.e.m. **p<0.01, ***p<0.001, ns = not significant (unpaired, two-tailed, Student’s *t*-test).

### ACLY associates with F-actin

Many of the known regulators of macropinocytosis (e.g. V-ATPase, RAC1, RAS, SDC1, SLC4A7 and PI3K) have been shown to localize to the plasma membrane (7). Therefore, as an initial step towards defining the mechanistic basis for the regulatory role of ACLY in macropinocytosis, we examined its subcellular distribution using immunofluorescence microscopy. Consistent with prior reports, we found that ACLY displayed a prominent nucleocytosolic localization (22) (Fig. 2A). In addition, the staining pattern revealed a previously uncharacterized spatial distribution of ACLY, namely its localization to the cell periphery. The specificity of the staining was validated using ACLY WT and KO cells (Fig. 2B). To further characterize the nature of the observed peripheral ACLY staining pattern, membrane ruffles were labeled with phalloidin (which selectively binds F-actin, an obligatory constituent of membrane ruffles) (6). Confocal microscopy analysis revealed the presence of ACLY in phalloidin-labelled membrane ruffles in multiple cell lines (Fig. 2C). The lack of apparent membrane targeting sequences on ACLY and the high degree of spatial overlap between ACLY and cortical F-actin (Fig. 2C) raised the possibility that ACLY is associated with the actin cytoskeleton. To test this idea, detergent fractionation was used to extract soluble, cytosolic proteins and lipid-based organelles, whilst retaining the F-actin cytoskeleton in the insoluble fraction. Immunoblot analysis of lysates from detergent extracted cells demonstrated the presence of ACLY in the detergent insoluble fraction in multiple cell lines (Fig. 2D and Fig. S2A). Loss of α-tubulin from the detergent insoluble fraction was used to assess extraction efficiency and fraction purity. Confocal immunofluorescence analysis of ACLY in detergent extracted cells confirmed the presence of a detergent-resistant pool of ACLY that colocalized with F-actin (Fig. 2E and Fig. S2B, C). Importantly, this staining pattern was lost in detergent-extracted cells following treatment with the F-actin depolymerizing agent latrunculin A, indicating that the retention of ACLY in the detergent insoluble fraction is mediated by its interaction with F-actin (Fig. 2E and Fig. S2B, C). Consistent with this interpretation, using biotinylated phalloidin to pulldown F-actin from cell lysates, we found that ACLY was associated with F-actin (Fig. 2F, G). Known F-actin binding proteins (ARP3 and ACTN1) and non-F-actin binding proteins (α-tubulin and PCNA) served as positive and negative controls for affinity purification, respectively. Ectopically expressed, catalytically inactive ACLY(H760A) retained the ability to co-localize with F-actin, indicating that this association was not dependent on the enzymatic activity of ACLY (Fig. S2D, E). In addition, the colocalization of ACLY with F-actin was also displayed in WT RAS-expressing HeLa cells, demonstrating that the ability of ACLY to associate with F-actin was not dependent on oncogenic RAS-mediated signaling (Fig. S2F, G). Collectively, these findings identify a previously unappreciated subcellular localization for ACLY that is mediated by its association with F-actin.

**Figure 2.**
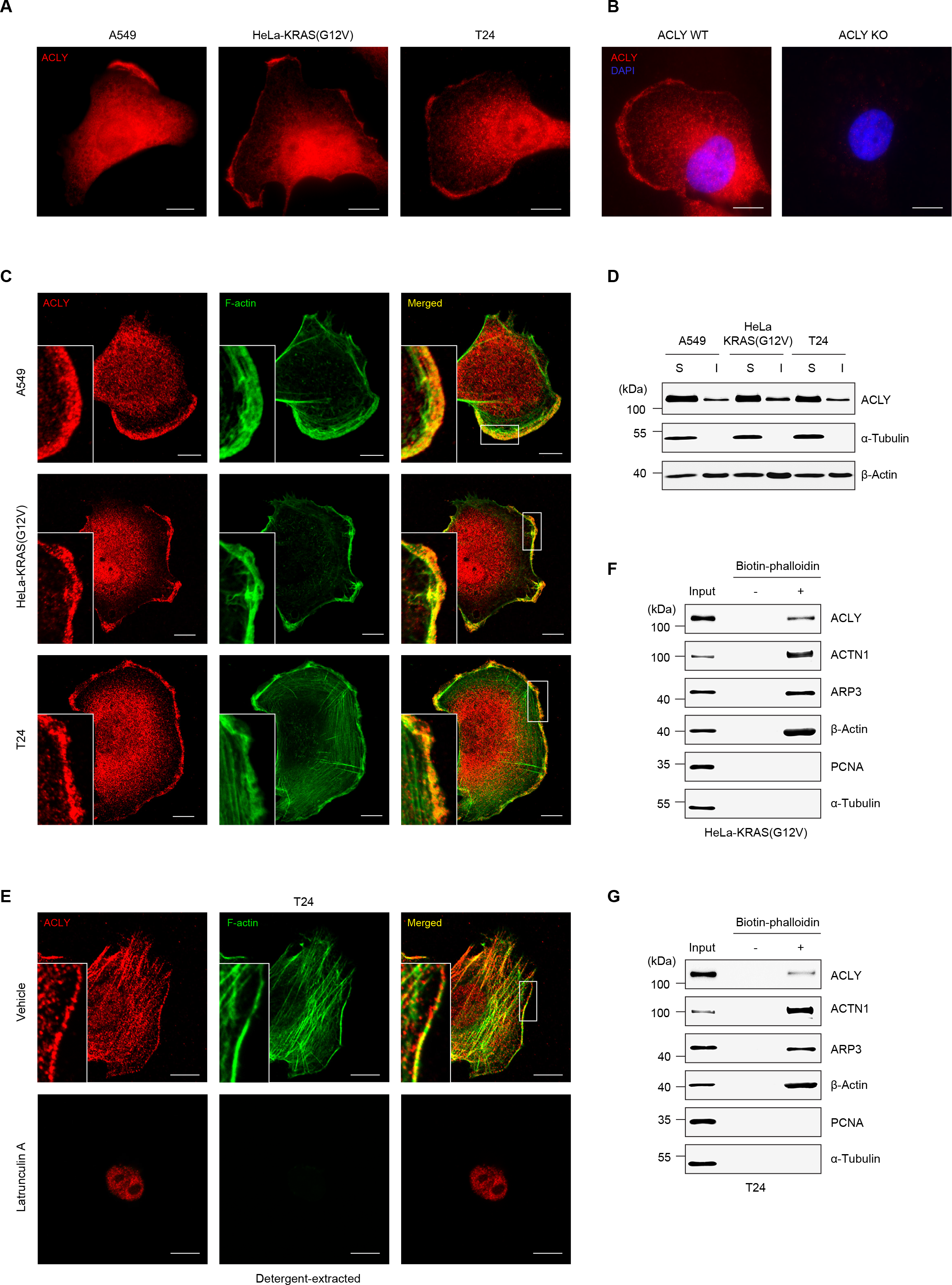
ACLY associates with F-actin. **(A)** Fluorescence micrographs (epifluorescence microscopy) of cells immunostained for ACLY showing nucleocytosolic and peripheral labelling of ACLY. **(B)** Fluorescence micrographs (epifluorescence microscopy) of ACLY WT and KO T24 cells immunostained for ACLY showing antibody specificity. **(C)** Fluorescence micrographs (confocal microscopy) of cells immunostained for ACLY showing colocalization between ACLY and F-actin (phalloidin). **(D)** Immunoblot analysis of ACLY in detergent soluble (S) and insoluble (I) fractions in detergent-extracted cells. α-tubulin was used to assess fraction purity and β-actin was used as a loading control. **(E)** Fluorescence micrographs (confocal microscopy) of ACLY immunostaining and co-labeled with phalloidin in detergent extracted T24 cells treated with latrunculin A (1 hour). **(F)** Immunoblots of biotinylated phalloidin F-actin pulldown in HeLa-KRAS(G12V) and **(G)** T24. ARP3 and ACTN1 (known actin binding proteins) and α- tubulin and PCNA (non-actin binding proteins) were used as positive and negative controls, respectively. Scale bar for all images = 10 μm. All images and immunoblots are representative.

### Capping protein-dependent actin remodeling regulates macropinocytosis

Next, we sought to investigate the mechanistic link between the cytoskeletal-association of ACLY and the regulation of macropinocytosis. To this end, we employed a proximity-dependent biotin identification (BioID) proteomic screen. To capture both short- and long-range proximal proteins, we generated BirA-ACLY fusion proteins with different sized linkers (16 or 46 amino acids) (Fig. S3A). Importantly, we confirmed that the localization of these fusion proteins mirrored that of endogenous ACLY, in terms of F-actin association, and production of comparable total levels of biotinylation, relative to BirA control (Fig. S3B-D). In order to enrich for cytoskeletal-associated proteins, the screen was performed by affinity purifying biotinylated proteins from detergent-extracted cells expressing BirA or BirA-ACLY. In addition to previously reported ACLY interaction partners such as GSK3β, PKA, NDPK and SEC16A (20, 23), several heretofore undocumented F-actin binding proteins were among the screen hits (Fig. 3A, Table S1). Furthermore, gene ontology analysis revealed a specific enrichment of actin-dependent cellular components, consistent with actin cytoskeleton association (Fig. S3E).

**Figure 3.**
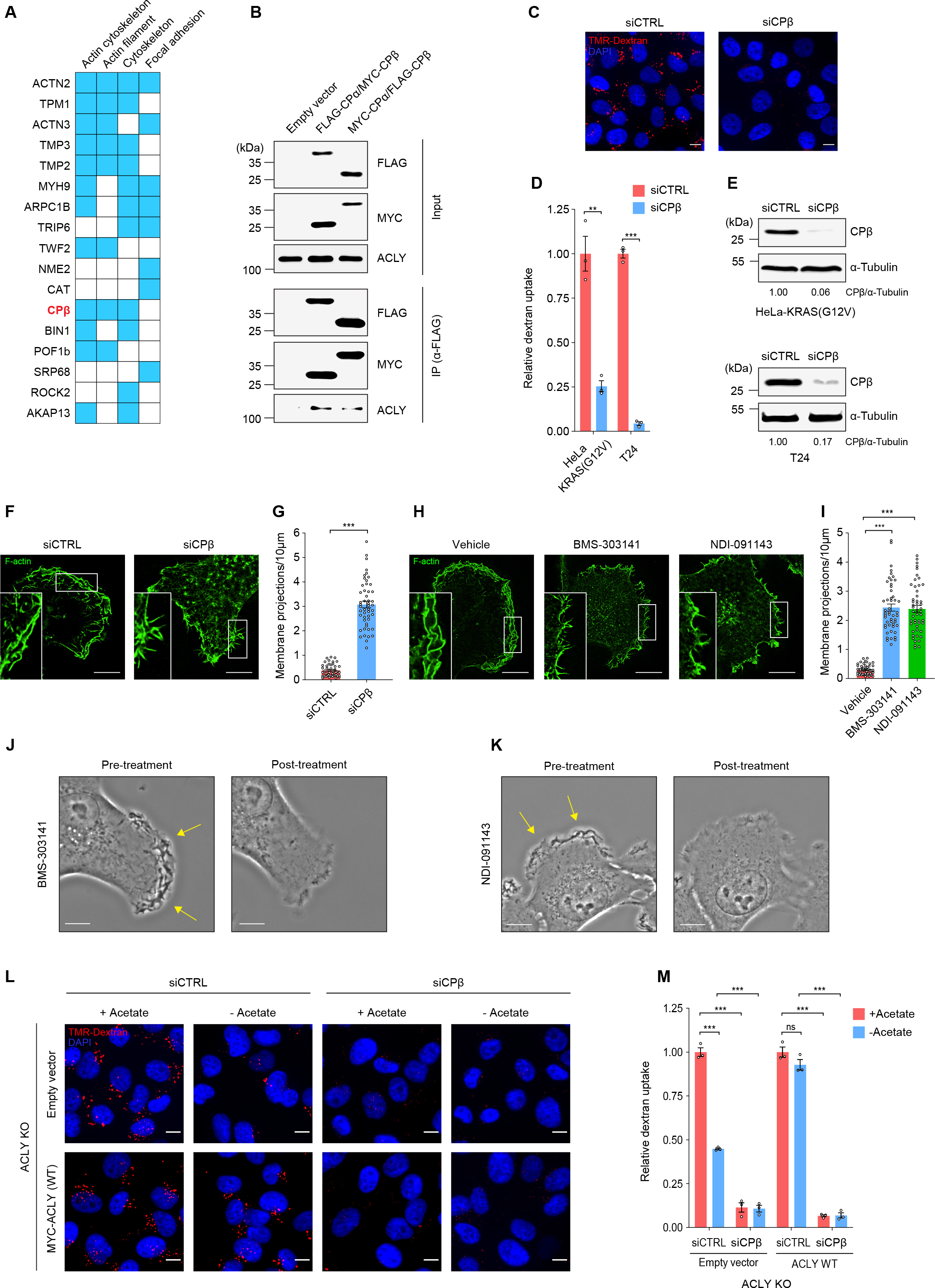
Actin capping protein controls actin dynamics and regulates macropinocytosis. (**A**) Gene ontology terms associated with BioID screen showing top actin-associated proteins among BirA-ACLY hits grouped by cellular component (Enrichr). Blue boxes indicate assignment to corresponding cellular component. **(B)** Co-immunoprecipitation (anti-FLAG) of endogenous ACLY with CP in HeLa-KRAS(G12V) cells. **(C)** Fluorescence micrographs of TMR-dextran uptake in T24 cells and **(D)** quantification in CPβ-depleted cell lines (72 hour knockdown). **(E)** Immunoblots showing CPβknockdown in HeLa-KRAS(G12V) and T24 cells (from dextran uptake analysis in D). α-tubulin was used as a loading control. **(F, G)** Fluorescence micrographs of F-actin (phalloidin) staining and quantification of membrane projections in T24 cells treated with siCPβ (72 hours) or **(H**, **I)** ACLY inhibitors (1 hour). The number of membrane projections were counted from 50 cells (each dot represents one cell) over 3 independent experiments and normalized to plasma membrane perimeter (per 10 μm). **(J)** Live phase contrast imaging of T24 cells treated with ACLY inhibitors (30 minutes), BMS-303141 and **(K)** NDI-091143. Yellow arrows indicate regions of active membrane ruffling. The same cells were imaged pre-treatment (vehicle) and post-treatment (inhibitor). **(L)** Fluorescence micrographs of TMR-dextran uptake and **(M)** quantification in CPβ-depleted (72 hour knockdown) ACLY KO T24 cells reconstituted with empty vector or WT ACLY in the presence or absence of sodium acetate. Scale bar for all images = 10 μm. All images and immunoblots are representative. For TMR-dextran uptake assays, at least 500 cells were counted per biological replicate (n=3). Data represented as mean ± s.e.m. **p<0.01, ***p<0.001, ns = not significant (unpaired, two-tailed, Student’s *t*-test).

In order to prioritize candidate proteins, we cross-referenced the BioID screen hits with genes from a genome-wide siRNA screen for macropinocytosis regulators previously performed by our group (24). Heterodimeric actin capping protein (CP) subunit beta (CPβ, encoded by *CAPZB*) was among the top hits from the siRNA screen (positive regulator of macropinocytosis) and was also detected in the BioID screen. Therefore, CPβ was selected as a candidate for further investigation. In line with BioID identification, we found that immunoprecipitates of ectopically expressed heterodimeric CP (achieved by endogenous co-expression of the CPα and CPβ subunits) contained endogenous ACLY, providing further evidence for their spatial proximity (Fig. 3B). CP, which binds to the ends of actin filaments and terminates polymerization, controls the architecture of the actin cytoskeleton by fine-tuning filament length and branching and regulating monomeric G-actin availability (25). Dysregulated F-actin capping, either by unrestrained capping (which causes attenuated polymerization) or insufficient capping (which leads to unrestricted polymerization and G-actin depletion) results in impairment in the cell’s ability to generate F-actin-driven protrusive membrane forces, a critical cytoskeletal remodeling determinant of membrane ruffles (25). In accordance with this postulated role, knockdown of CPβ potently inhibited dextran uptake (Fig. 3C-E and Fig. S4A, B). This was accompanied by the loss the lamellar architecture of membrane ruffles and the pronounced accumulation of thin projections of plasma membrane, a morphological feature previously shown to be induced by loss of CP function (26) (Fig. 3F, G). Significantly, similar phenotypic changes in membrane architecture were observed following pharmacological inhibition of ACLY, indicating that ACLY activity may regulate macropinocytosis through modulation of F-actin capping dynamics that are required for membrane ruffle formation (Fig. 3H-K). Consistent with this idea, we found that neither reconstitution of ACLY, nor acetate supplementation, were sufficient to restore dextran uptake in ACLY KO cells in the setting of CPβ depletion (Fig. 3L, M and Fig. S4C). These complementation experiments place CPβ as an essential downstream mediator of ACLY- and acetyl-CoA-dependent regulation of macropinocytosis.

### ACLY controls CP acetylation

We next set out to determine the mechanistic basis for the functional interdependence between acetyl-CoA and CP in the context of regulation of macropinocytosis. In agreement with data from proteome-wide acetylation screens demonstrating that CPβ is acetylated at multiple lysine residues (27–29), we detected acetylated CPβ using an antibody that recognizes acetylated-lysine (AcK) residues (Fig. 4A). The abundance of acetylated CPβ was significantly increased following treatment with pan-histone deacetylase inhibitors (TSA and NAM), confirming specificity of the anti-AcK antibody (Fig. 4A). The requirement of ACLY for the maintenance of CPβ acetylation was established by demonstrating that treatment of cells with ACLY inhibitors led to a significant reduction in the levels of acetylated CPβ (Fig. 4B). Furthermore, disruption of the F-actin cytoskeleton network using latrunculin A, which results in the preferential loss of F-actin-associated ACLY (Fig. 2E), caused a significant reduction in CPβ acetylation, but had no effect on histone acetylation (Fig. 4C). The dependence of CPβ acetylation on cytoskeletal-associated ACLY suggests that the anchoring of ACLY to cortical F-actin could serve as a mechanism for the tight spatiotemporal regulation of actin cytoskeletal dynamics through the reversible modulation of protein acetylation. Using mass spectrometry analysis to verify CPβ acetylation status and identify acetylation sites, we detected several tryptic peptides with lysine acetylation corresponding to positions K78, K199, K235, K237 and K254 (Fig. 4D). Importantly, the latter four residues are located at the C-terminus of CPβ (Fig. 4E) which has been shown to be essential for its binding to F-actin (30–32).

**Figure 4.**
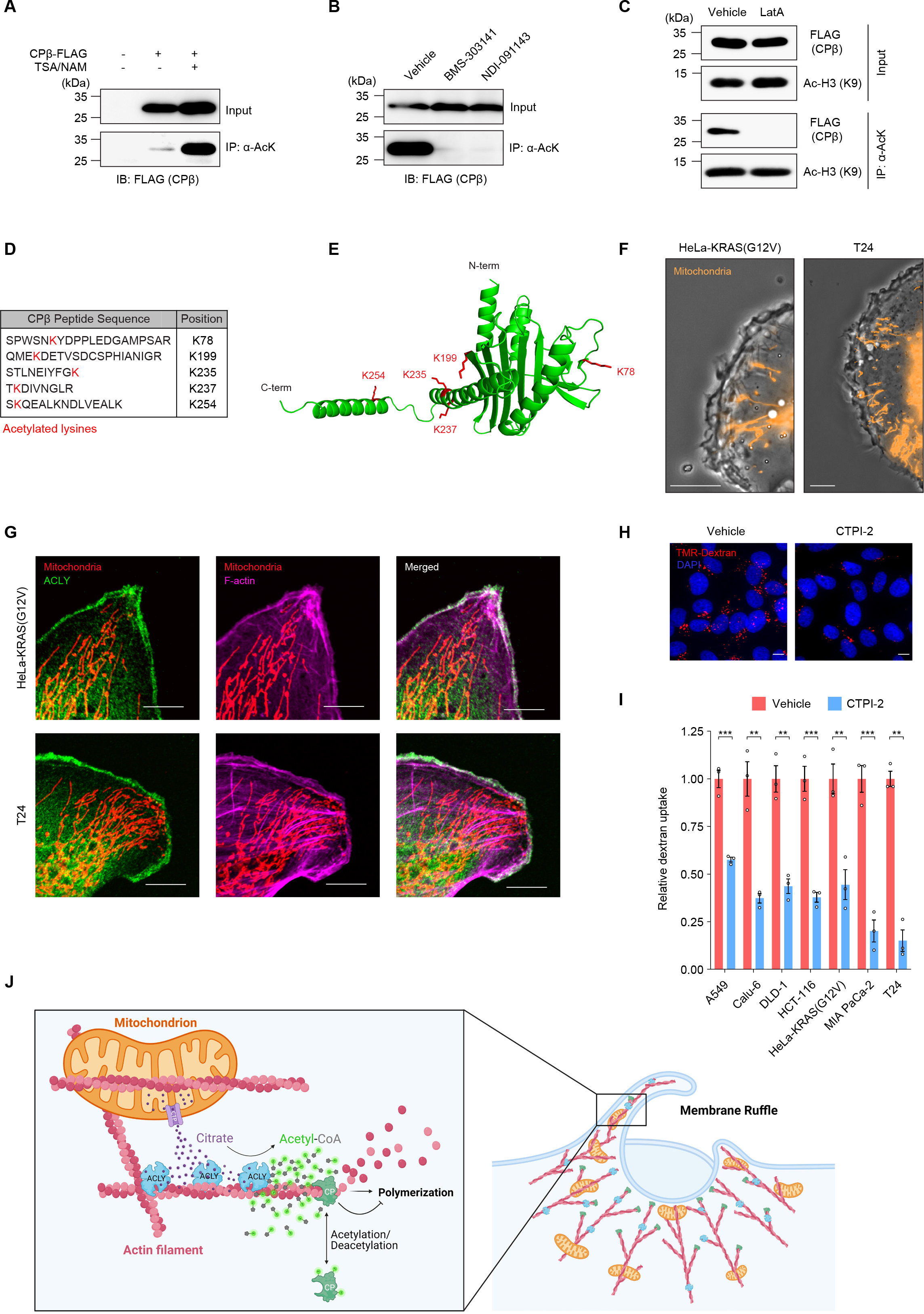
ACLY modulates actin capping protein acetylation. **(A)** Immunoblot analysis of acetylated CPβ-FLAG immunoprecipitated with anti-acetylated lysine antibody (anti-AcK) from HeLa-KRAS(G12V) cells treated with pan-histone deacetylase inhibitors trichostatin A (TSA) and nicotinamide (NAM, 5 hours), **(B)** ACLY inhibitors BMS-303141 and NDI-091143 (2 hours) or **(C)** latrunculin A (Lat A, 1 hour). Ac-H3 (K9) is acetylated histone H3 (lysine 9). **(D)** Tryptic peptide sequences from mass spectrometry post-translation modification (PTM) analysis showing the positions of acetylated lysine residues (red) in CPβ-FLAG immunoprecipitated (anti-FLAG) from HeLa-KRAS(G12V) cells. **(E)** Structure of human CPβ protein (PyMOL) showing positions of acetylated lysine residues (red) identified from mass spectrometry PTM analysis in D. (**F)** Images of live phase contrast/fluorescence microscopy of HeLa-KRAS(G12V) and T24 expressing Mito-DsRed (mitochondrial probe) showing mitochondria in proximity to membrane ruffles. **(G)** Fluorescence micrographs (confocal microscopy) of HeLa-KRAS(G12V) and T24 cells expressing Mito-DsRed immunostained for ACLY and co-labeled with phalloidin (F-actin), showing mitochondria in proximity to membrane ruffle-associated ACLY. **(H)** Fluorescence micrographs of TMR-dextran uptake in T24 cells and **(I)** quantification in cell lines treated with a citrate transport protein (CTP) inhibitor, CTPI-2 (1.5 hours). **(J)** Schematic of model showing mitochondria infiltration into actin-rich membrane ruffles. CTP-dependent export of citrate from membrane ruffle-localized mitochondria is used by actin-associated ACLY to locally generate acetyl-CoA that is required for the spatially defined acetylation of capping protein (CP). Acetylation regulates the F-actin localization of CP, which controls actin polymerization and membrane ruffle formation. Scale bar for all images = 10 μm. All images and immunoblots are representative. For TMR-dextran uptake assays, at least 500 cells were counted per biological replicate (n=3). Data represented as mean ± s.e.m. **p<0.01, ***p<0.001 (unpaired, two-tailed, Student’s *t*-test).

A prerequisite for the proposed role of F-actin-associated ACLY in the localized production of acetyl-CoA is the channeling of its substrates (mitochondrial-derived ATP and citrate) into the vicinity of cortical ACLY. In fact, mitochondrial positioning at the leading edge of the cell has been shown to play an important role in supporting the ATP-dependent cytoskeleton rearrangements that drive cell migration (33, 34). In support of this idea, we observed by live cell imaging and immunofluorescence analysis that mitochondria traffic to the cell periphery towards areas of membrane ruffling (Fig. 4F, Videos S1-2) where they are found in close proximity to membrane ruffle-associated ACLY (Fig. 4G). Given that utilization of mitochondrial-derived citrate by cytosolic ACLY requires its export via the mitochondrial citrate transporter protein (CTP, encoded by *SLC25A1*), we tested whether blocking mitochondrial citrate export affects macropinocytosis. In accordance with a requirement of macropinocytosis for mitochondrial-derived citrate, pharmacological inhibition of CTP significantly impaired dextran uptake (Fig. 4H, I). These findings support a model wherein submembranous channeling of mitochondrial-derived citrate into cytoskeleton-localized ACLY might serve as a mechanism by which the acetylation-dependent actin dynamics that are required for membrane ruffling and macropinocytosis can be spatially fine-tuned (Fig. 4J).

## Discussion

Macropinocytosis is now recognized as a central metabolic adaptive process that is utilized by tumor cells to scavenge extracellular macromolecules under nutrient limiting conditions. In this study we identify a previously unidentified step in the regulatory circuitry that controls macropinocytosis involving the localized production of acetyl-CoA by actin-associated ACLY. Our findings provide new insights into the molecular underpinning of macropinocytosis, and underscore the importance of understanding spatial determinants of the cellular machinery that drive this process.

Dynamic acetylation, which engenders a means of rapid and reversible modulation of protein activity and localization, is required for regulation and execution of many fundamental biological processes including cell signaling, transcription, cell cycle regulation and cytoskeleton organization (35). Consistent with the essential role of acetyl-CoA as the obligatory acetyl donor for acetylation reactions (36), the intracellular abundance of acetyl-CoA is a major determinant of global acetylation levels (18, 22). However, there is mounting evidence that local acetyl-CoA availability within defined cellular compartments contributes to the spatiotemporal control of acetylation (37–41). These observations suggest that regulation of the subcellular localization of acetyl-CoA producing enzymes can be essential for specifying the spatiotemporal dynamics of protein acetylation.

In this study we identify CPβ (the β subunit of heterodimeric CP) as a target for location specific ACLY-dependent acetylation. CP has been shown to be required for the formation of many actin driven cellular structures, including membrane ruffles, a function that is dependent on its ability to bind to the ends of actin filaments (25). There is evidence to suggest that several basic residues that lie within the F-actin-binding interface, which spans the C-terminal portion of the protein, promote the interaction of CP with F-actin (30–32). Of relevance, several of the CPβ acetylation sites identified in this study are located within this interface. Given that acetylation results in lysine charge neutralization, it is plausible that the acetylation of one or more of the lysine residues within the F-actin binding interface might affect the electrostatic interactions between CPβ and F-actin. Indeed, acetylation of CP has been suggested to stimulate actin dynamics in phenylephrine-stimulated hypertrophying myocytes by promoting the dissociation of CP from F-actin (42). The identity of the lysine acetyltransferases (KATs) and lysine deacetylases (KDACs) that mediate the reversible acetylation of CP remains to be determined. Of note, several KDACs and KATs have been shown to localize to the actin cytoskeleton (42–44) suggesting that the control of actin assembly by dynamic acetylation can be spatially regulated at multiple levels.

While the association of ACLY with the actin cytoskeleton identified in the present study has been investigated specifically in the context of the F-actin remodeling machinery that regulates membrane ruffling and macropinocytosis, its functional relevance might extend to other ACLY-mediated cellular processes. For example, nuclear F-actin has been shown to play an essential role in promoting the repair of DNA double strand breaks (DSBs) (45, 46). In fact, many DNA repair factors have been shown to interact with F-actin, which has been suggested to contribute to their stabilization at sites of damage (47). Notably, nuclear ACLY has been implicated in the regulation of DNA repair through the localized production of acetyl-CoA, which promotes the acetylation of histones at DSBs, and subsequently stimulates the recruitment of DNA repair factors (41). These findings suggest that the capacity of ACLY to interact with the actin cytoskeleton could be also critically important for the spatial orchestration of the DNA damage response.

Genetic and pharmacological blockade of ACLY has been shown to inhibit tumor growth in various in vivo cancer models (17, 20, 48). Thus far, the protumorigenic function of ACLY has largely been attributed to acetyl-CoA-dependent lipid biosynthesis and histone acetylation (13). By implicating ACLY in the regulation of macropinocytosis, our study suggests a new modality by which ACLY might contribute to the metabolic fitness of tumor cells, namely by sustaining their capacity to support anaplerotic pathways via the macropinocytic uptake of extracellular proteins. Additionally, by virtue of playing a role in submembranous F-actin assembly, a process required for the formation of membrane ruffles that are known to generate the driving force for cell movement (49), ACLY likely serves as an essential component of the regulatory network that controls cancer cell migration and invasion. Future studies examining the mechanisms underlying the association of ACLY with F-actin, and their impact on spatially defined F-actin dynamics, will provide new insights into the links between metabolic reprogramming and actin-dependent processes that drive cell transformation.

## Methods

### Reagents and DNA constructs

All reagents used were analytical grade. TMR–Dextran (70 kDa) was purchased from Fina Biosolutions. BMS-303141 (SML0784), CTPI-2 (ENA018104423), puromycin (P8833), doxycycline (D9891), biotin (B4639) trichostatin A (TSA, T8552) and nicotinamide (NAM, N0636) were purchased from Sigma. NDI-091143 was purchased from Aobious (AOB17806). Latrunculin A was purchased from Cayman Chemical (10010630). Protease Inhibitor Cocktail was purchased from Roche (11697498001). The following primary antibodies were used in this study: anti-α-tubulin (clone B-5-1-2 - Sigma, T5168); anti-acetylated histone H3 (K9) (clone C5B11- Cell Signaling, 9649); anti-acetylated-lysine (Ac-K2-100) MultiMab (Cell Signaling, 9814); anti- ACLY (Sigma, HPA022434); anti-ACTN1 (Abclonal, A1160); anti-ARP3 (clone FMS338 - Abcam, ab49671); anti-β-actin (clone 13E5 - Cell Signaling, 4970); anti-CPβ (Bethyl Laboratories, A304-734A); anti-histone H3 (clone D1H2 - Cell Signaling, 4499); anti-MYC (clone 9B11, Cell Signaling, 2276); anti-PCNA (clone PC10 - Cell Signaling, 2586). For immunoblotting, the following secondary antibodies were used: Alexa Fluor 680 goat anti-mouse (Invitrogen, A21058); IRDye 800CW goat anti-rabbit (Li-Cor, 926-32211), HRP-conjugated goat anti-rabbit (Cell Signaling, 7074,); HRP-conjugated goat anti-mouse (Cell Signaling, 7076). For immunofluorescence, Alexa Flour 488-conjugated goat anti-rabbit (Invitrogen, A11029), Alexa Flour 555-conjugated goat anti-rabbit (Invitrogen, A21429) or Alexa Flour 555-conjugated goat anti-mouse (Invitrogen, A21424) secondary antibodies were used. The following siRNAs were purchased from Dharmacon (Horizon):

Non-targeting siRNA Pool #2 (D-001206-14): UAAGGCUAUGAAGAGAUAC;
AUGUAUUGGCCUGUAUUAG; AUGAACGUGAAUUGCUCAA;
UGGUUUACAUGUCGACUAA.
siCPβ SMARTPool (M-011990-01): GAAGUACGCUGAACGAGAU;
GGAGUGAUCCUCAUAAAGA; GAGACAAGGUGGUGGGAAA;
CACCAUGGAGUAACAAGUA.

pTRIPZ-KRAS(G12V) was constructed by cloning human, codon optimized KRAS(G12V) with an N-terminal T7 tag into pTRIPZ (Dharmacon) using *AgeI* and *MluI* to simultaneously remove RFP. Human ACLY(H760A) was generated by site-directed mutagenesis using the pCMV6-ACLY(MYC-FLAG) vector (Origene, RC200508). Human CPα (CAPZA1, cloneID 3047937) and CPβ (CAPZB, clone ID 6715709) were purchased from Dharmacon and inserted into pcDNA3 (Invitrogen) with N-terminal MYC or FLAG tags. BirA-ACLY constructs were generated by inserting human ACLY into pcDNA3.1 mycBioID vector (N-terminal MYC-BirA, Addgene, plasmid 35700) with a 16 or 46 amino acid linkers between ACLY and BirA. ACLY lentivirus constructs were generated by inserting ACLY variants with an N-terminal MYC tag into pLVX-IRES-Puro (Clontech, 632183). All DNA constructs were verified by Sanger sequencing.

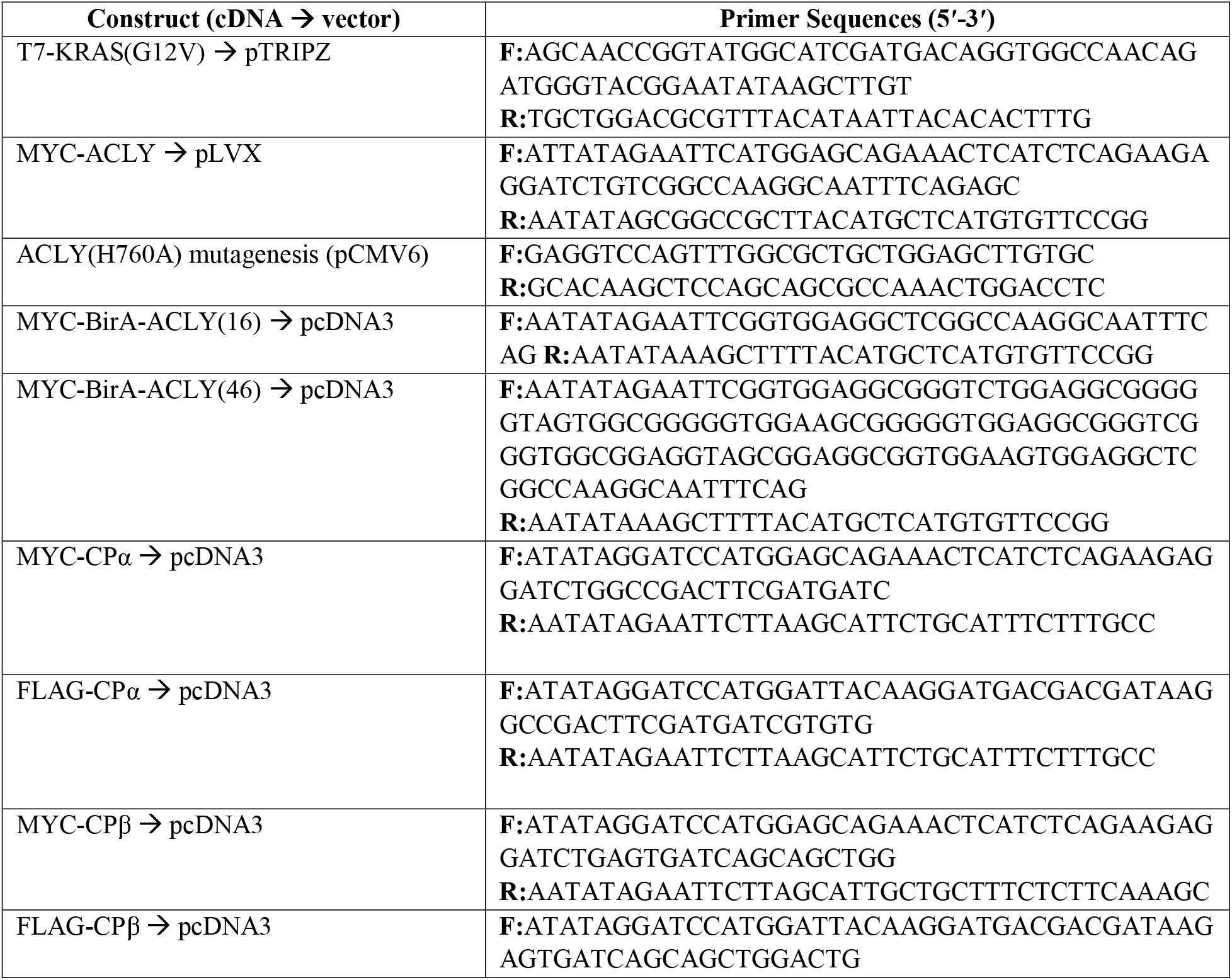
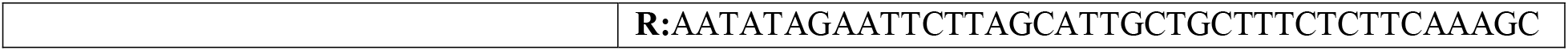

### Cell culture, treatment conditions and generation of stable cell lines

All cell lines (HeLa, A549, Calu-6, DLD-1, HCT-116, MIA PaCa-2, T24 and 293T) were purchased from the American Type Culture Collection and routinely tested negative for mycoplasma. Cells were maintained at 37°C with 5% CO_2_ in high glucose DMEM (Gibco, 11965-092) supplemented 1% penicillin/streptomycin (Gibco, 15140-122) and 10% FBS (Gibco, 10438-026). Inducible HeLa-KRAS(G12V) cells were generated by transducing HeLa cells with pTRIPZ-T7-KRAS(G12V) lentiviral particles followed by selection with puromycin (2 μg/mL) for 72 hours. HeLa-KRAS(G12V) cells were maintained in 10% tetracycline-free FBS (Takara Bio, 631101) and KRAS(G12V) expression was induced using doxycycline (250 ng/mL for 48 hours). Reconstitution of ACLY in ACLY KO T24 cells was performed by pooling all four KO clones in equal ratios and infecting with pLVX-IRES-Puro-(MYC-ACLY) lentiviral particles. 48 hours after transduction, cells were selected with puromycin (2 μg/mL) for 72 hour. Cells stably expressing the Mito-DsRed mitochondrial probe were generated by infecting cells with pLV-mitoDsRed (Addgene, plasmid 44386) lentivirus particles with puromycin (2 μg/mL) selection for 72 hour. Lentivirus particles were packaged in 293T cells by co-transfecting carrier plasmids with psPAX (Addgene, plasmid 12260) and psPAX2 (Addgene, plasmid 12260). Virus-containing supernatant was concentrated using Amicon Ultra-15 100kDa MWCO centrifugal filters (Millipore, UFC910024). For siRNA knockdown experiments, siRNA (20nM) was complexed with Lipofectamine RNAiMax Transfection Reagent (Thermo Fisher Scientifc, 13778075) in OptiMEM (Thermo Fisher Scientifc, 31985062) according to the manufacturer’s instructions. Transfection complexes were added to cells in complete media for 6 hours before replacing media. For DNA transfections, DNA was complexed with X-tremeGENE 9 DNA Transfection Reagent (Sigma, 6365779001) in OptiMEM according to the manufacturers’ instructions. Unless otherwise indicated, drug treatment conditions were maintained the same throughout experimental assays. For BMS-303141 (5μM) and NDI-091143 (10μM), cells were treated with inhibitor for 1 hour in serum-free DMEM after 4 hours of serum-starvation. For CTPI-2 (50μM), cells were treated for 90 minutes in serum-free DMEM after 4 hours of serum-starvation. For anti-AcK immunoprecipitation, cells were treated with TSA (2.5μM) and NAM (5mM) for 5 hours in serum-free DMEM. For acetate withdrawal dextran uptake experiments, ACLY WT and KO cells were cultured in complete DMEM containing 10% dialyzed FBS with or without sodium acetate (1mM) for 5 hours followed by 3 hours serum-starvation with or without sodium acetate (1mM). Latrunculin A (200nM) was used for 1 hour in serum-free media.

### CRISPR/Cas9 genome editing

A sgRNA construct targeting exon 2 of the human ACLY gene was made by inserting annealed, phosphorylated oligonucleotides (5′-CACCGGAATCGGTTCAAGTATGCTC-3′, 5′-AAACGAGCATACTTGAACCGATTCC-3′) into pSpCas9(BB)-2A-Puro (PX459, Addgene, plasmid 48139). The sgRNA construct was then transfected (2 μg) into T24 cells using Lipofectamine 3000 (Invitrogen, 100022050) according to the manufacturers’ instructions. Cells were selected with puromycin (2 μg/mL) for 48 hour then single cell sorted by FACS into 96 well plates. Non-edited clones (i.e. without mutations in ACLY) were derived from the puromycin-selected population and used as ACLY WT controls. ACLY WT and KO cells were maintained in 1mM acetate. Clones were verified by Sanger sequencing of PCR products spanning the sgRNA target site generated using the following primers: 5′-CCTTCTGACCAGCTTCTCTCTCC-3′ (F) and 5′-GGCATCACCAACAAACCAATGGC-3′ (R).

### Macropinosome visualization and quantification

Macropinocytic index was quantified using dextran uptake as previously described (3). Briefly, cells were seeded onto glass coverslips and allowed to adhere for 48-72 hours. Cells were serum-starved for 4 hours and treated as indicated before incubation with TMR-dextran (70kDa) diluted in serum-free media (1mg/mL) containing indicated treatments for 30 minutes at 37°C. Cells were then fixed with 3.7% formaldehyde solution for 15 minutes at room temperature and stained with DAPI (1μg/mL). Coverslips were mounted using anti-fade mounting media (DKAO, S3023). Images were captured using a Ti2 Eclipse Epifluorescence Microscope (Nikon). Image analysis and quantification was performed in ImageJ (v1.53q) using raw data files as previously described (3).

### Live cell imaging

Cells were seeded into glass bottom 6-well plates in FluoroBrite DMEM (Gibco, A1896701) and allowed to adhere for 48 hours. Cells were serum starved for 4 hours in a pre-equilibrated, humidified chamber (37°C, 5% CO_2_) of a Ti2 Eclipse Epifluorescence Microscope (Nikon). Phase contrast and fluorescence images were captured every 10 seconds for 30-60 minutes using 60X or 100X oil objectives. For inhibitor experiments, drugs were added directly to the media (i.e. without replacing media) to minimize temperature and pH changes. Image brightness and contrast adjustments were performed in ImageJ (v1.53q) using raw data files.

### Immunofluorescence

Cells were seeded onto glass coverslips in 24-well plates. 48-72 hours after seeding, cells were fixed with 3.7% formaldehyde solution for 15 minutes at room temperature followed by two washes in PBS. Cells were then incubated in blocking solution (5% normal goat serum in 0.1% Triton X-100/PBS) for 30 minutes at room temperature and incubated with primary antibodies (1:250, anti-ACLY and 1:500, anti-MYC) diluted in blocking solution overnight at 4°C. Cells were then washed in PBS, incubated with secondary antibodies diluted 1:1000 in blocking solution, washed in PBS and stained with Alexa Fluor 488-conjugated phalloidin (Invitrogen, A12379) or Alexa Fluor 647-conjugated phalloidin (Abcam, ab176759) for 1 hour in 1% BSA/PBS. Coverslips were mounted using anti-fade mounting media (DKAO, S3023). For detergent extraction, prior to fixation, cells were incubated on ice for 5 minutes in freshly prepared, ice cold extraction buffer: 10 mM HEPES (pH 7.4), 50 mM NaCl, 2.5mM MgCl, 300 mM sucrose, 0.5% Triton X-100, 1 mM SOV, 1 mM NaF and Protease Inhibitor Cocktail. Cells were then washed in extraction buffer (without Triton X-100) and fixed as above. Loss of α- tubulin from the detergent soluble fraction, which occurs through cold-induced depolymerzation of microtubules, was used as a control to assess fraction purity. Images were captured using a LSM510 META Confocal Microscope (Zeiss) or a Ti2 Eclipse Epifluorescence Microscope (Nikon). Brightness and contrast adjustments were performed in ImageJ (v1.53q) using raw data files. See ‘Reagents and DNA constructs’ for list of primary and secondary antibodies used.

### Immunoprecipitation and immunoblotting

For immunoprecipitation experiments, cells were seeded into 15cm plates and transfected with indicated constructs using X-tremeGENE 9 (Sigma, 6365779001) according to the manufacturer’s instructions. Cells were washed with ice cold PBS before being scrapped, pelleted and lysed in lysis buffer: 50 mM Tris (pH 7.4), 250 mM NaCl, 1 mM EDTA, 1mM MgCl, 0.5% NP40, 1 mM DTT, 5 mM NAM, 2.5 μM TSA, 1 mM SOV, 1 mM NaF and Protease Inhibitor Cocktail. Lysates were then sonicated, cleared by centrifugation and incubated overnight with anti-acetylated lysine antibody (1:100) or for 1 hour with 50μL of pre-washed α- FLAG M2 affinity gel agarose slurry (Sigma, A2220) on a rotator at 4°C. For acetylated-lysine IP, antibody capture was performed with Protein A Dynabeads (Invitrogen, 10001D) for 2 hours on a rotator at 4°C. Beads were washed five times in lysis buffer then proteins eluted by boiling in sample buffer (40mM Tris pH 6.8, 1% SDS, 5% β-mercaptoethanol, 7.5% glycerol) for 10 minutes. For whole cell lysates, cells were washed in PBS, scraped in sample buffer then lysates sonicated and boiled for 10 minutes. Cell lysates were resolved by SDS-PAGE and electrotransferred onto nitrocellulose or PVDF membranes, which were then blocked using 5% skim milk in TBS-T (0.1% Tween-20) for 1 hour at room temperature. Membranes were then incubated with primary antibodies diluted in 5% BSA/TBS-T for either 1 hour at room temperature or overnight at 4°C with shaking. Following primary antibody incubations, membranes were washed in TBS-T then incubated for 1 hour at room temperature with secondary antibodies diluted 1:10,000 in blocking buffer. For detection of biotinylated proteins, HRP-conjugated streptavidin (ThermoFisher Scientific, SA10001) diluted 1:5000 in 5% BSA/TBS-T was used. For HRP detection, blots were developed with SuperSignal West Pico PLUS Chemiluminescent Substrate (Thermo Scientific, 34579) or SuperSignal West Femto Maximum Sensitivity Substrate (Thermo Scientific, 34095). Imaging of membranes was performed using an ImageQuant800 Analyzer (GE Healthcare) or an Odyssey Scanner (Li-Cor). For quantification of protein levels from immunoblots, Odyssey Software (Li-Cor) was used to measure fluorescence intensity of individual bands. See ‘Reagents and DNA constructs’ for list of primary and secondary antibodies used.

### Proteomics sample preparation and mass spectrometry

For BioID screening, HeLa-KRAS(G12V) cells were grown in 15cm dishes and transfected with pcDNA3.1-BirA-Myc (2 μg + 8 μg pcDNA3 as a carrier plasmid) or pcDNA3.1-BirA-Myc-ACLY (10 μg) using X-tremeGENE 9 (Sigma, 6365779001). At the time of transfection, cells were treated with doxycycline (250 ng/mL) and biotin (50 μM). 30h after transfection, cells were washed with PBS then incubated with ice cold extraction buffer (freshly prepared) for 5 minutes on ice: 10 mM HEPES (pH 7.4), 50 mM NaCl, 2.5mM MgCl, 300 mM sucrose, 0.5% Triton X-100, 1 mM SOV, 1 mM NaF and Protease Inhibitor Cocktail. Extracted cells were washed with extraction buffer (without Trixon X-100) then scraped and lysed in RIPA buffer: 50 mM Tris-HCl (pH 7.4), 250 mM NaCl, 5 mM EDTA, 0.5% DOC, 0.1% SDS, 1% NP40, 1 mM SOV, 1
mM NaF and Protease Inhibitor Cocktail. Lysates were incubated for 30 minutes at 4°C on a rotator then cleared by centrifugation (10,000 x g). Cleared lysates were incubated with 300 μL pre-washed NeutrAvidin Agarose slurry (Thermo Scientific, 29200) for 1 hour at 4°C on a rotator. Beads were washed twice in 5 mL of high salt RIPA containing 500mM NaCl (5 minutes at 4°C on a rotator) then twice with RIPA (5 minutes at 4°C on a rotator). Beads were then resuspended in sample buffer (40mM Tris pH 6.8, 1% SDS, 5% β-mercaptoethanol, 7.5% glycerol), passed through a column (Chromotek, sct-50) to remove beads and submitted for mass spectrometry analysis. For CPβ-FLAG PTM analysis, HeLa-KRAS(G12V) cells were grown in 15cm dishes and transfected with pcDNA3-FLAG (15 μg) or pcDNA3-CPβ-FLAG (7.5 μg) and pcDNA3-CPα-MYC (7.5 μg) using X-tremeGENE 9 (Sigma, 6365779001). At the time of transfection, cells were treated with doxycycline (250 ng/mL). 30h after transfection, cells were washed in PBS, collected, pelleted (2000 x g) then resuspended in lysis buffer: 50 mM Tris (pH 7.4), 250 mM NaCl, 1 mM EDTA, 1mM MgCl, 0.5% NP40, 5% glycerol, 1 mM DTT, 2 μM TSA, 5 mM NAM, 1 mM SOV, 1 mM NaF and Protease Inhibitor Cocktail. Lysates were incubated for 30 minutes on ice, sonicated and cleared by centrifugation (10,000 x g). Cleared lysates were incubated with 300 μL of pre-washed α-FLAG M2 affinity gel agarose slurry (Sigma, A2220) and incubated for 1 hour at 4°C on a rotator, then beads were washed three times in lysis buffer. Immunoprecipitated proteins were eluted with 0.5 mg/mL (diluted in lysis buffer) 3X FLAG peptide (Sigma, F4799) for 30 minutes on a rotator. Eluates were then centrifuged and supernatants passed through a column (Chromotek, sct-50) to remove residual beads before concentrating using 3 kDa MWCO Amicon Ultra-0.5 centrifugal filters (Millipore, UFC5003). Concentrated eluates were submitted for mass spectrometry analysis. Samples were prepared for mass spectrometry analysis by reducing with DTT (2 μL of 0.2M) for 1 hour at 57 °C and alkylatation with iodoacetamide (2 μL of 0.5M) for 45 minutes at room temperature in the dark. Proteins were resolved on a NuPAGE 4-12% Bis-Tris gel (Life Technologies) then excised and gel slices dehydrated with methanol (100mM) ammonium bicarbonate (100mM) mixture before digestion with 200ng of trypsin (Promega) overnight at room temperature. Peptides were extracted by incubating gel slices with R2 POROS beads (Thermo Scientific) followed by acidification with 0.1% trifluoroacetic acid (TFA) and desalted using Sep-Pak C18 solid-phase extraction (Waters). Peptides were eluted with 40% acetonitrile in 0.5% acetic acid then 80% acetonitrile in 0.5% acetic and concentrated using a SpeedVac and stored at −80°C. Peptides were gradient eluted from the column directly into the Q Exactive mass spectrometer (Thermo Scientific) using a 1 hour gradient with solvent A (2% acetonitrile, 0.5% acetic acid) and solvent B (80% acetonitrile, 0.5% acetic acid). High resolution full MS spectra were acquired with a resolution of 45,000, an AGC target of 3×10^6^, a maximum ion time of 45 milliseconds, and scan range of 400-1500 m/z. Following each full MS, twenty data-dependent high resolution higher-energy C-trap dissociation (HCD) MS/MS spectra were acquired. All MS/MS spectra were collected using the following instrument parameters: resolution of 15,000, AGC target of 1×10^5^, maximum ion time of 120 milliseconds, 1 microscan, 2 m/z isolation window, fixed first mass of 150 m/z, and normalized collision energy (NCE) of 27. MS/MS spectra were searched against a Uniprot Human database using Proteome Discoverer 1.4 (for BioID screen) or against human CPβ protein using Byonic (for PTM analysis).

### Biotinylated phalloidin pulldown

72 hour after seeding into 15cm dishes, cells were washed twice with ice cold PBS, scraped and pelleted. Cell pellets were suspended in lysis buffer: 50 mM Tris (pH 7.4), 50 mM KCl, 150 mM NaCl, 1 mM EDTA, 1mM MgCl2, 0.5% Triton X-100, 1mM ATP, 1 mM DTT, 1 mM SOV, 1 mM NaF and Protease Inhibitor Cocktail. Lysates were incubated for 15 minutes on ice then cleared by centrifugation (3000 x g). Supernatant was collected and pre-cleared with protein G agarose beads (Pierce, 20397) for 15 minutes at 4°C on a rotator. Beads were removed by centrifugation (3000 x g for 1 minute) and lysates split equally and incubated with either methanol (vehicle) or 1 μM biotinylated phalloidin (Biotium, 00028) for 2 hour at 4°C on a rotator. Lysates were then incubated with 30μL of pre-washed NeutrAvidin agarose resin (Thermo Scientific, 29200) for 1 hour at 4°C on a rotator. Beads were washed 5X in lysis buffer then resuspended in sample buffer (40mM Tris pH 6.8, 1% SDS, 5% β-mercaptoethanol, 7.5% glycerol), boiled for 10 minutes then subjected to SDS-PAGE and immunoblotting.

### Statistical analysis

Unless otherwise stated, data in all graphs are represented as mean ± standard error of the mean (s.e.m.) of n ≥ 3 biological replicates. Two-tailed, unpaired Student’s *t*-tests (equal variance) was used for statistical comparisons between groups (GraphPad Prism v9.2.0). P-values and number of samples analyzed for each experiment is indicated in figure legends.

## Data Availability

Mass spectrometry proteomic data will be made available by depositing in the appropriate database.

## Acknowledgements

This work was supported by funding from the National Institutes of Health (NIH)/National Cancer Institute (NCI) Grant CA210263 (D.B.S). J.P. was supported by a fellowship from the National Health and Medical Research Council (NHMRC) CJ Martin Biomedical Fellowship (APP1106545). The mass spectrometric experiments were supported in part by NYU Langone Health and the Laura and Isaac Perlmutter Cancer Center support grant P30CA016087 from the NCI and by the NIH Shared Instrumentation Grant 1S10OD010582-01A1 for the purchase of an Orbitrap Fusion Lumos Tribrid mass spectrometer. All original data and reagents are available from the corresponding authors upon request.

