## Supplementary figures and images for "Cytoskeletal association of ATP citrate lyase controls the mechanodynamics of macropinocytosis"

### Supp. Figures

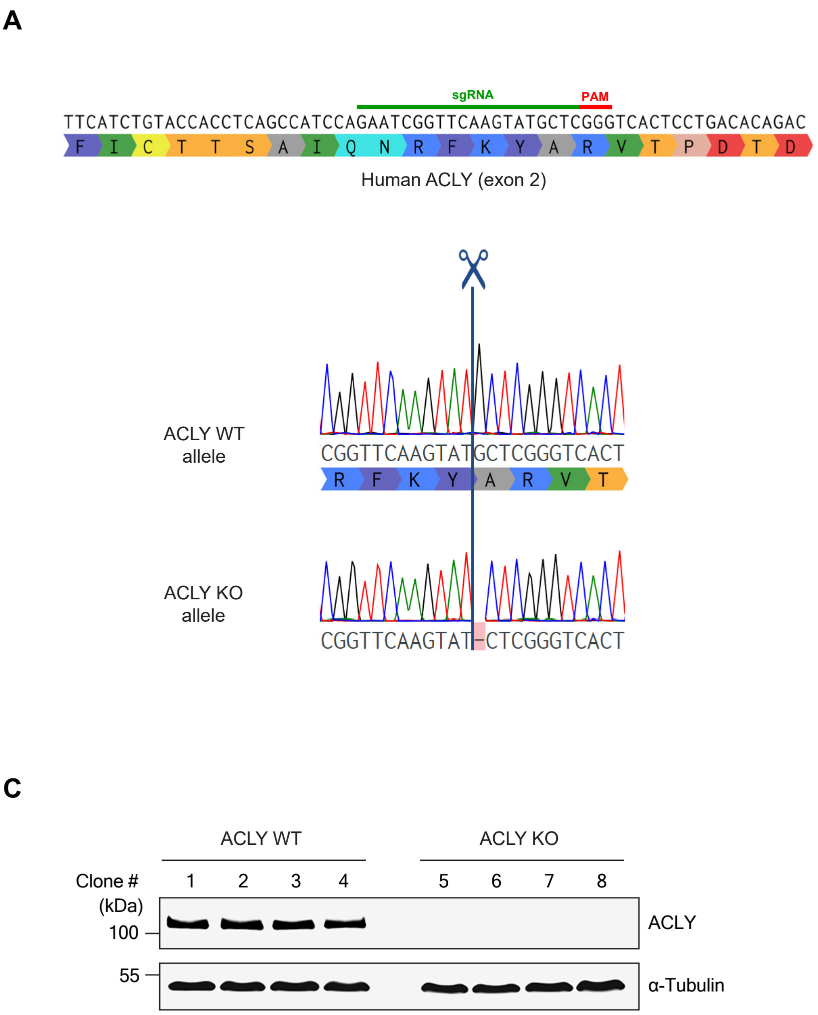


Figure S1, Puccini et al.

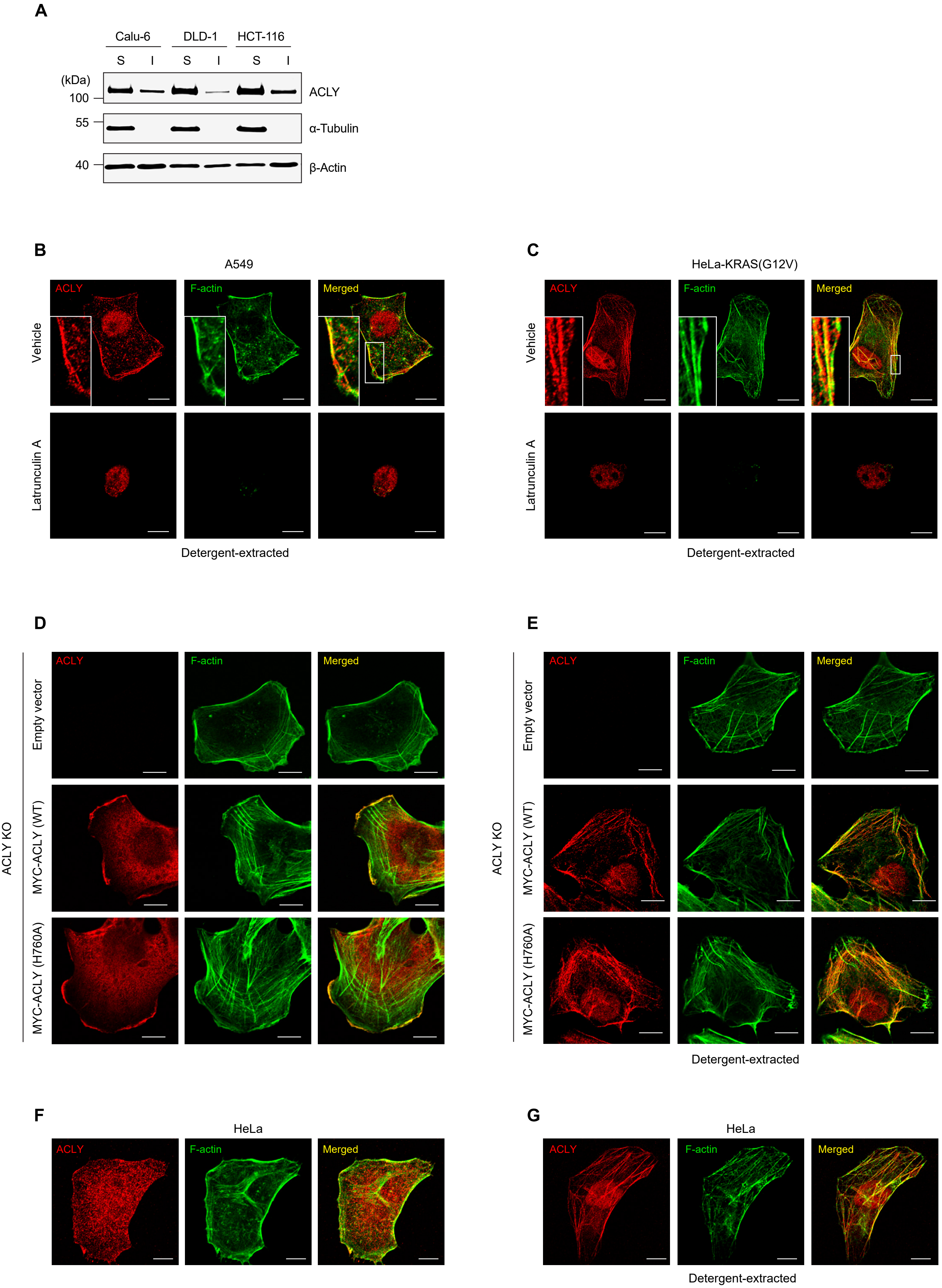


Figure S2, Puccini et al.

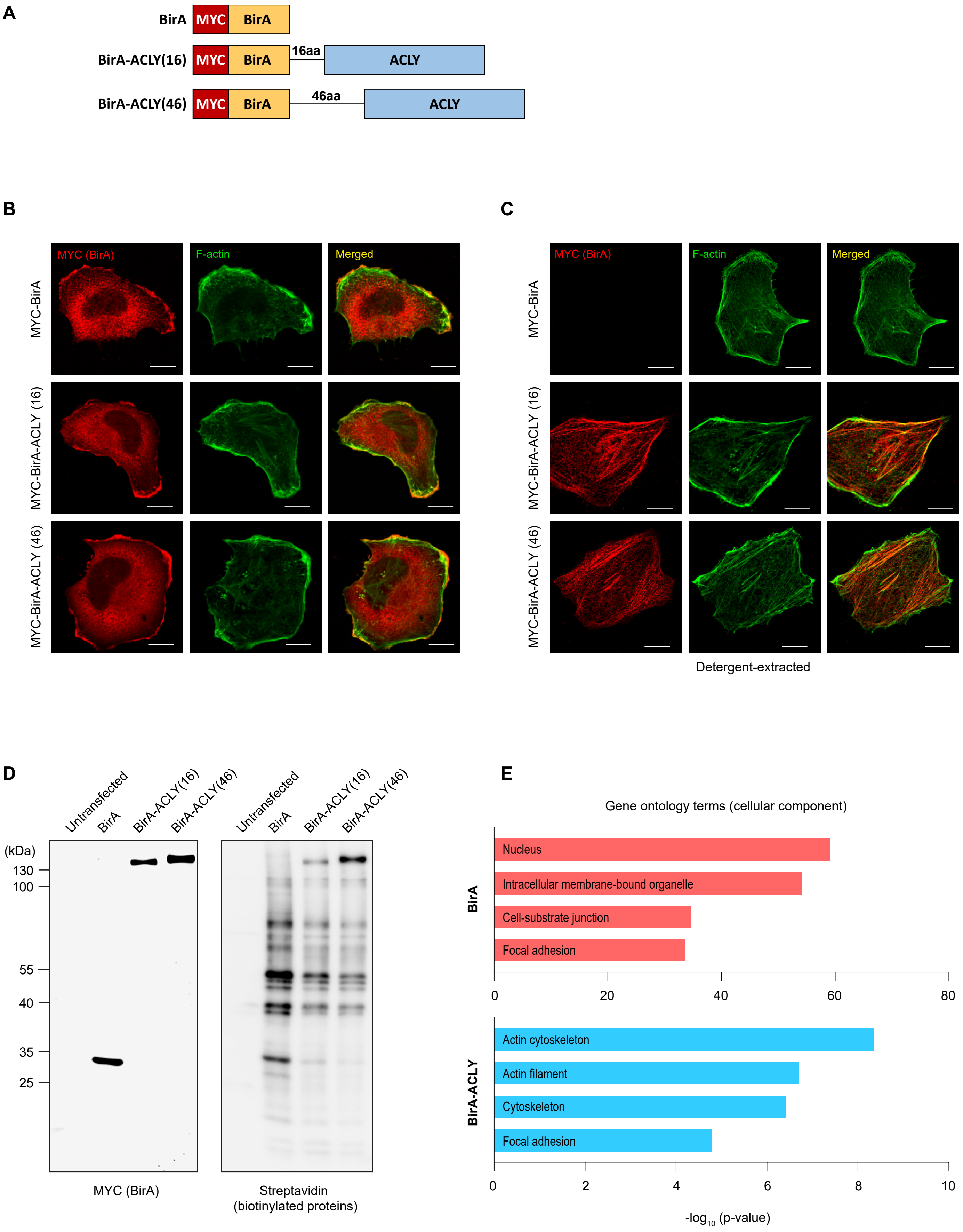


Figure S3, Puccini et al.

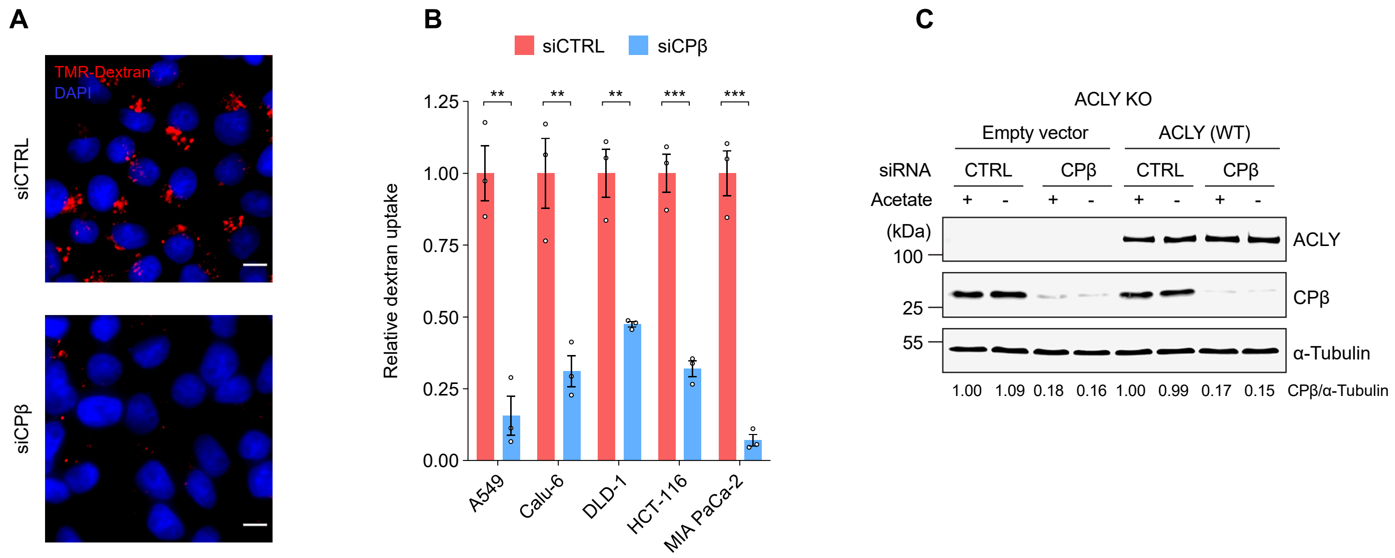


Figure S4, Puccini et al.
