## Supplementary material for "Cytoskeletal association of ATP citrate lyase controls the mechanodynamics of macropinocytosis": Supp. Figure Legends

**Figure S1. Generation of ACLY KO clones**. **(A)** Schematic showing genomic locus and protein

sequence of human ACLY (exon 2) with sgRNA targeting site and representative Sanger

sequencing traces of ACLY wild type (WT) and ACLY knockout (KO) T24 clones. **(B)**

Immunofluorescence and **(C)** immunoblot analysis of endogenous ACLY expression in ACLY

WT and ACLY KO T24 clones. α-tubulin was used as a loading control. Scale bar for all images

= 10 μm. All images and immunoblots are representative.

**Figure S2. ACLY associates with F-actin**. **(A)** Immunoblot analysis of ACLY in detergent

soluble (S) and insoluble (I) fractions in detergent-extracted cells. α-tubulin was used to assess

fraction purity and β-actin was used as a loading control**. (B)** Fluorescence micrographs (confocal

microscopy) of cells immunostained for ACLY and co-labeled with phalloidin (F-actin) in

detergent-extracted A549 and **(C)** HeLa-KRAS(G12V) cells treated with latrunculin A (1 hour).

**(D)** Fluorescence micrographs (confocal microscopy) of cells immunostained for ACLY and colabeled

with phalloidin in non-extracted and **(E)** detergent-extracted ACLY KO T24 cells

reconstituted with empty vector, WT ACLY-MYC or H760A (catalytically inactive mutant)

ACLY-MYC. **(F)** Fluorescence micrographs (confocal microscopy) of HeLa cells (expressing WT

RAS) immunostained for ACLY and co-labelled with phalloidin in non-extracted and **(G)**

detergent-extracted cells. Scale bar for all images = 10 μm. All images and immunoblots are

representative.

**Figure S3. BioID proteomic screen for ACLY proximal proteins. (A)** Schematic showing the

BirA-ACLY fusion proteins used for BioID screen with 16 and 46 amino acid linkers between

BirA and ACLY. **(B)** Fluorescence micrographs (confocal microscopy) of non-extracted and **(C)**

detergent-extracted HeLa-KRAS(G12V) cells expressing MYC-BirA or MYC-BirA-ACLY

fusion proteins immunostained for MYC and co-labeled with phalloidin (F-actin). **(D)** Immunoblot

analysis of MYC-BirA and MYC-BirA-ACLY expression and biotinylated proteins from

streptavidin affinity purification from BioID screen. **(E)** Gene ontology terms associated with

ACLY BioID screen hits according to cellular compartment (Enrichr). Scale bar for all images =

10 μm. All images and immunoblots are representative.

**Figure S4. Actin capping protein is required for macropinocytosis. (A)** Fluorescence

micrographs of TMR-dextran uptake in MIA PaCa-2 and **(B)** quantification in cell lines transfected

with siCPβ (72 hours). **(C)** Immunoblots showing CPβ knockdown and ACLY expression in

ACLY KO T24 cells reconstituted with empty vector or WT ACLY (from dextran uptake analysis

in Fig. 3M). α-tubulin was used as a loading control. For TMR-dextran uptake assays, at least 500

cells were counted per biological replicate (n=3). Scale bar for all images = 10 μm. All images and

immunoblots are representative. Data represented as mean ± s.e.m. **p<0.01, ***p<0.001

(unpaired, two-tailed, Student’s *t*-test).
